# Giantin is required for intracellular N-terminal processing of type I procollagen

**DOI:** 10.1101/2020.05.25.115279

**Authors:** Nicola L. Stevenson, J. M. Bergen Dylan, Chrissy L. Hammond, David J. Stephens

## Abstract

Knockout of the golgin giantin leads to skeletal and craniofacial defects driven by poorly studied changes in glycosylation and extracellular matrix deposition. Here, we sought to determine how giantin impacts the production of healthy bone tissue by focussing on the main protein component of the osteoid, type I collagen. Giantin mutant zebrafish accumulate multiple spontaneous fractures in their caudal fin, suggesting their bones may be more brittle. Inducing new experimental fractures revealed defects in the mineralisation of newly deposited collagen as well as diminished procollagen reporter expression in mutant fish. Analysis of giantin knockout cells expressing a GFP-tagged procollagen showed that procollagen trafficking is independent of giantin. However, our data show that intracellular N-propeptide processing of pro-α1(I) is defective in the absence of giantin. These data demonstrate a conserved role for giantin in collagen biosynthesis and extracellular matrix assembly. Our work also provides evidence of a giantin-dependent pathway for intracellular procollagen processing.

## Introduction

The golgins are a family of coiled-coil domain proteins that extend out from the surface of the Golgi apparatus to tether transport vesicles and other Golgi membranes (Munro, 2011). The largest member of this family, giantin, is a tail-anchored membrane protein with a predicted 37 cytosolic coiled-coil domains (Linstedt and Hauri, 1993; Seelig et al., 1994). These structural features are key attributes for a membrane tether, however, to date no tethering function for giantin has been identified. Indeed, giantin loss does not block anterograde transport (Lan et al., 2016; Stevenson et al., 2017) and may in fact accelerate it (Koreishi et al., 2013). Most studies also agree that giantin is not essential to maintain Golgi morphology (Koreishi et al., 2013; Lan et al., 2016; Puthenveedu and Linstedt, 2001; Stevenson et al., 2017), although it may inhibit lateral tethering between cisternae (Satoh et al., 2019; Stevenson et al., 2017). Discrepancies between these studies are likely due to variation in levels of depletion (Stevenson et al., 2017), in genetic compensation (Stevenson et al., 2017), and/or functional redundancy with other golgins (Wong and Munro, 2014).

The most consistent observation from published work is that giantin is required to regulate glycosylation (Kikukawa et al., 1990; Koreishi et al., 2013; Lan et al., 2016; Petrosyan et al., 2014; Stevenson et al., 2017) and extracellular matrix (ECM) formation (Katayama et al., 2018; Kikukawa and Suzuki, 1992; Lan et al., 2016). Highly selective defects in *O*-glycosylation have been reported following knockout (KO) of the *GOLGB1* gene encoding giantin in cells (Stevenson et al., 2017), zebrafish (Stevenson et al., 2017), and mice (Lan et al., 2016). Enzyme distribution (Petrosyan et al., 2014) and surface glycosylation patterns (Koreishi et al., 2013) are more generally affected following siRNA depletion. The secretion of ECM proteoglycans and collagen can also be affected (Katayama et al., 2018; Kikukawa et al., 1990). The primary phenotype shared by all *GOLGB1* KO animal models is the abnormal development of craniofacial structures, whilst species specific phenotypes include short limbs in rats (Katayama et al., 2011) and ectopic mineralisation of soft tissues in zebrafish (Stevenson et al., 2017). Giantin is therefore important for skeletal development and defects in ECM quality likely underly all these phenotypes.

In light of these observations, we hypothesised that giantin may regulate secretion of the primary protein component of skeletal ECM, fibrillar type I collagen. In mammals, this is predominantly built from heterotrimeric molecules composed of two pro-α1(I) chains (encoded by the *COL1A1* gene) and one pro-α2(I) chain (encoded by *COL1A2*) chain. Structurally, each chain is made up of a helical domain, composed of Gly-X-Y repeats, flanked by globular N- and C-propeptide domains (Canty and Kadler, 2005). These chains are co-translationally translocated into the endoplasmic reticulum (ER) lumen where they are post-translationally modified prior to folding into a right-handed triple helical molecule. Trimeric procollagen is exported from the ER and transits the Golgi before being secreted from the cell and assembled into fibrils in the ECM (Canty and Kadler, 2005; Canty et al., 2004).

Prior to fibrillogenesis, the N- and C-propeptide domains of procollagen are cleaved to promote correct alignment and polymerisation. Removal of the C-propeptide is particularly critical as this induces self-assembly of collagen into fibrils (Hulmes et al., 1989; Kadler et al., 1987; Kadler et al., 1990; Miyahara et al., 1984; Miyahara et al., 1982). Retention of the N-propeptide on the other hand does not preclude fibril assembly but can affect fibril morphology (Bornstein et al., 2002; Hulmes et al., 1989; Romanic et al., 1992). C-terminal processing is carried out by BMP-1/tolloid-like family metalloproteinases (Kessler et al., 1996) while ADAMTS2, −3 and −14 cleave the N-propeptide (Bekhouche and Colige, 2015; Colige et al., 1997; Colige et al., 2002; Fernandes et al., 2001). Meprin α and meprin β have also been implicated in procollagen processing (Broder et al., 2013).

In this study, by using a combination of fin fracture assays in *golgb1* mutant zebrafish and biochemical assays in giantin KO cells, we demonstrate that giantin function is required to facilitate normal fracture repair and for intracellular N-terminal processing of type I procollagen.

## Results

### Homozygous golgb1 mutant fish have a higher incidence of fracture

To begin our investigation into the role of giantin in the deposition of skeletal ECM we first examined our previously published homozygous (HOM) *golgb1^X3078/X3078^* mutant zebrafish line for bone defects (Stevenson et al., 2017). Focussing on the experimentally tractable caudal fin (Bergen et al., 2019), we observed an unusually high number of naturally occurring fractures in the hemirays of homozygous (HOM) individuals compared to wild type (WT) and heterozygote (HET) siblings. This was seen both in terms of the number of injured fish and the number of fractures per individual. Indeed at 7 months old, 76% of HOM fish had acquired at least one fracture compared to just 33% of WT and 27% of HET fish (Figure 1A). The mean number of fractures per individual was 0.4 for WT and HETs and 1.8 for mutants. Interestingly, four or more fractures were detected in 12% of *golgb1^X3078/X3078^* mutants. This was never seen in WT or HET siblings. These were often in the proximal half of the fin and occurred either consecutively along one ray or adjacent to each other in parallel rays (Figure 1B). There was also one HOM individual carrying fractures that had fused together, indicating aberrant fracture repair with excessive calcification (Figure 1B).

**Figure 1.**
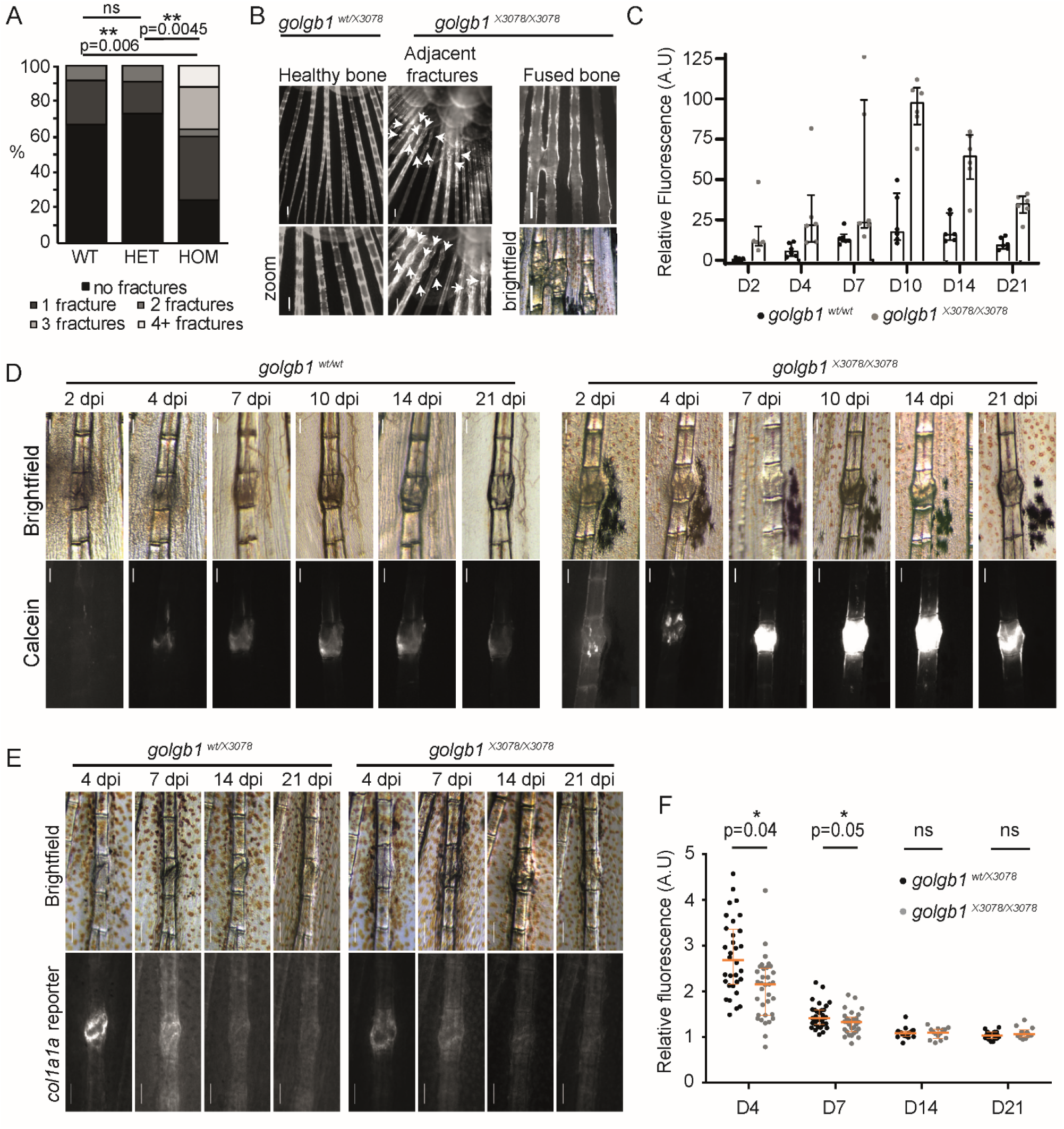
Fracture defects in *golgb1* mutant zebrafish. **A.** Quantification of the number of fractures naturally found in the caudal fin of WT and *golgb1^X3078^* heterozygous and homozygous mutant fish at 7 months old. Data show % of fish with x number of fractures (WT=12 fish, HET=11 fish, HOM=25 fish from 2 independent crosses). **B**. Fluorescence images of naturally occurring caudal fin defects in *golgb1^X3078/X3078^* fish stained with alizarin red. White arrows indicate fractures. Scale bar 200 μm. **C-D.** Quantification (C) and representative brightfield and fluorescent images (D) of experimentally induced fractures in *golgb1^wt/wt^ and golgb1^X3078/X3078^* caudal fins on different days post-injury (dpi). Bone was stained with calcein at each timepoint prior to imaging. (C) Calcein intensity in fractures was measured relative to that of healthy adjacent bones. Lower exposure images than those in D were used for quantification to avoid saturation. Each dot represents one fracture. Bars show median and interquartile range (2 fish per line quantified, each with 3 fractures). (D) Proximal end of bone is at the top of image. Scale bar 200 μm. **E.** Representative brightfield and fluorescent images of experimentally induced fractures in the caudal fin of *golgb1^wt/X3078^ and golgb1^X3078/X3078^* fish expressing a *col1a1a:GFP* promoter reporter at different timepoints. Proximal end of bone is at the top of image. Scale bar 200 μm. **F.** Quantification of *col1a1a:GFP* signal at the fracture site relative to an adjacent healthy bone. Each dot represents one fracture. Orange bars indicate median and interquartile range. At timepoints 4 and 7 d.p.f eleven fish per line were quantified. At timepoints 14 and 21 d.p.f n= 6 HETs and n=4 HOM fish quantified. All data was collected in a single experiment. All p values were calculated with the Mann-Whitney U test comparing the means for each fish, 3 fractures per fish.

### *golgb1* mutant fish show mineralisation defects during fracture repair

To look at fractures more closely, fractures were experimentally induced in caudal fin hemirays of mutant and WT fish as has been previously described (Geurtzen et al., 2014; Tomecka et al., 2019). Injured fish were stained with calcein at different timepoints to monitor calcification of newly formed bone matrix in the callus. Calcein labelling was first visible in WT fish four days post injury (dpi) whereas labelling was already apparent at the fracture site in HOM fish at 2 dpi (Figure 1C-D). In both cases labelling intensity continued to increase over time, peaking at 10 dpi. Calcein fluorescence was substantially greater in the fractures of HOM fish throughout the assay. This is most evident at 10 dpi when it was approximately 400% brighter in the mutants. Levels of calcein-accessible calcium are therefore elevated in the fractures of *golgb1^X3078/X3078^* fish. This shows that premature and enhanced calcification occurs in the mutant fish.

### Expression of the *col1a1* gene is reduced in *golgb1* mutant fractures

Mineralisation is dependent on the correct spatio-temporal deposition of a collagen matrix to ensure the right amount of calcification occurs at the right time (Michigami, 2019). We therefore looked at type I collagen expression in the repairing fractures. We crossed *golgb1^X3078/X3078^* fish with a *col1a1a:GFP* promoter reporter line and performed the same fracture experiments as above but without the calcein stain. Expression of *col1a1a:GFP* at the fracture site relative to healthy bone was highest at 4 dpi in both HET and HOM fractures (Figure 1E-F). Expression returned to normal levels by 14 dpi. Interestingly, *col1a1a:GFP* expression was lower in the mutant fractures compared to HETs at both 4 and 7 dpi when promoter activity was at its peak. This agrees with our previously published RNAseq data which shows a reduction in *COL1A1* mRNA in *GOLGB1* KO hTERT-RPE1 cells relative to WT (Stevenson et al., 2017). No obvious difference in callous width or alizarin red staining was observed (Supplemental Figure 1).

### Expression and secretion of type I collagen is elevated in giantin KO cells

To better understand the matrix defects observed above, we turned to our previously published *GOLGB1* KO hTERT-RPE1 cell line to study collagen secretion in a more tractable system (Stevenson et al., 2017). Our previously published RNAseq data shows that gene expression of *COL1A1* is reduced in giantin KO cells (Stevenson et al., 2017). In contrast, immunoblot analysis shows that KO cells contain higher levels of pro-α1(I) protein relative to WT cells (Figure 2A-B). While statistical testing of pooled data does not reveal a detectable difference here, comparison of individual experiments (colour-coded in Figure 2B) shows a consistent increase of pro-α1(I). Interestingly, higher procollagen levels are not due to protein retention since KO cells also secrete similar or higher amounts of pro-α1(I) compared to WT cells, both in absolute terms and relative to the intracellular pool (Figure 2C-D). There is also no evidence of collagen over-modification or ER dilation (Stevenson et al., 2017) as would be expected if the collagen were retained (Ishida et al., 2006). Immunofluorescence labelling of pro-α1(I) in non-permeabilised WT and KO cells verified that this excess of secreted collagen is incorporated into the extracellular matrix (Figure 2E).

**Figure 2.**
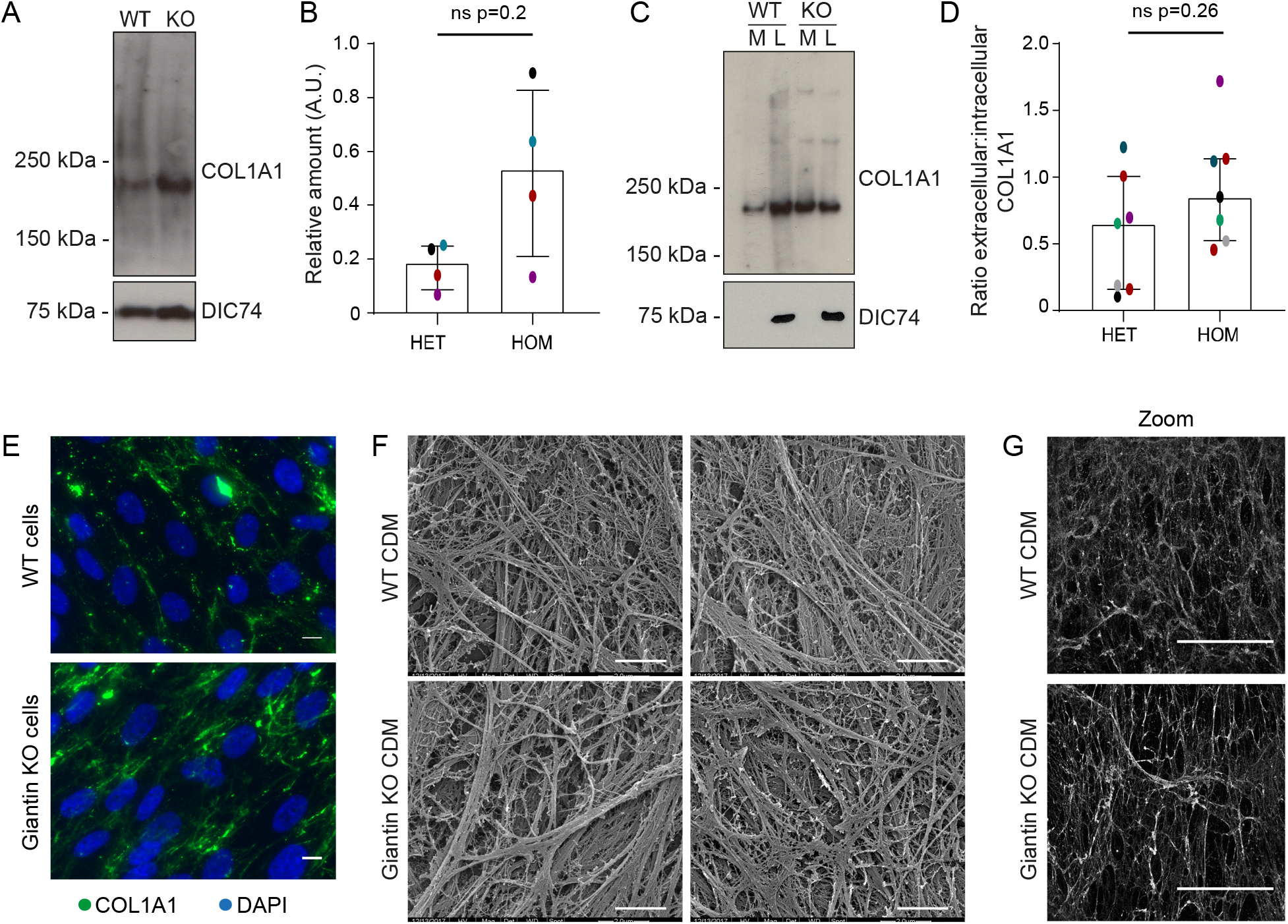
pro-α1(I) is more abundant in giantin KO cells. **A**. Western blot of pro-α1(I) (COL1A1) and dynein intermediate chain (DIC74, housekeeping) in cell lysates taken from WT and giantin KO cells. **B.** Densitometry of the semi-quantitative ECL western blots represented in (A). Dots show individual replicates and each independent experiment is colour-coded between cell lines. Bars depict median intensity of pro-α1(I) (COL1A1) normalised against DIC74 (n=4 biological replicates). Error bars = interquartile range. Statistical test Mann-Whitney **C.** Western blot of media (M) and cell lysate (L) fractions taken from WT and giantin KO cell cultures after 16 h incubation with serum free medium plus 50 μg/μl ascorbate. **D.** Ratio of extracellular vs intracellular levels of collagen as measured from secretion assays represented in (C). pro-α1(I) (COL1A1) levels measured by densitometry from semi-quantitative ECL blots and normalised against DIC74 before calculating ratios. Dots show individual replicates and each independent experiment is colour-coded between cell lines. Bars depicts median and interquartile range (n=7 biological replicates). Statistical test Mann-Whitney **E**. Maximum projections of widefield image stacks showing PFA fixed, unpermeabilised cells immunolabelled for endogenous pro-α1(I) (COL1A1, green). Nuclei are stained with DAPI (blue). Scale bars 10 μm. **F-G**. The cell derived matrix produced by WT and giantin KO cells imaged as (A) scanning electron micrograms, and (B) confocal z stacks of antibody labelled pro-α1(I) presented as maximum projections. Scale bars 2 μm (A) and 100 μm (B).

To determine whether the organisation of the matrix produced by KO cells is altered compared to WT, we stripped the cell layer from 7-day confluent cultures and imaged the cell derived matrix (CDM) both by scanning election microscopy (Figure 2F) and by confocal microscopy after immunolabelling for pro-α1(I) (Figure 2G). In each case, the giantin KO CDM often appeared to contain thicker fibres that were more intensely labelled for pro-α1(I).

### Procollagen trafficking through the early secretory pathway is unaffected by loss of giantin

We next tested whether altered procollagen synthesis and deposition is the result of changes to procollagen trafficking through the secretory pathway. To this end, we generated WT and giantin KO cell lines stably expressing low levels of proSBP-GFP-COL1A1 (GFP-COL1A1, (McCaughey et al., 2019)). This construct encodes pro-α1(I) with a GFP and SBP tag inserted upstream of the N-propeptide cleavage site (Figure 3A). The SBP tag allows controllable bulk release of the procollagen from the ER using the Retention Using Selective Hooks assay (RUSH, (Boncompain et al., 2012)). This assay relies on co-expression of a KDEL-streptavidin hook to bind the SBP and anchor the GFP-COL1A1 in the ER. The experimental addition of biotin then displaces the SBP, releasing the procollagen. The hook is co-expressed from a bicistronic vector with mCherry-sialyltransferase (mCh-ST) which labels the Golgi.

**Figure 3.**
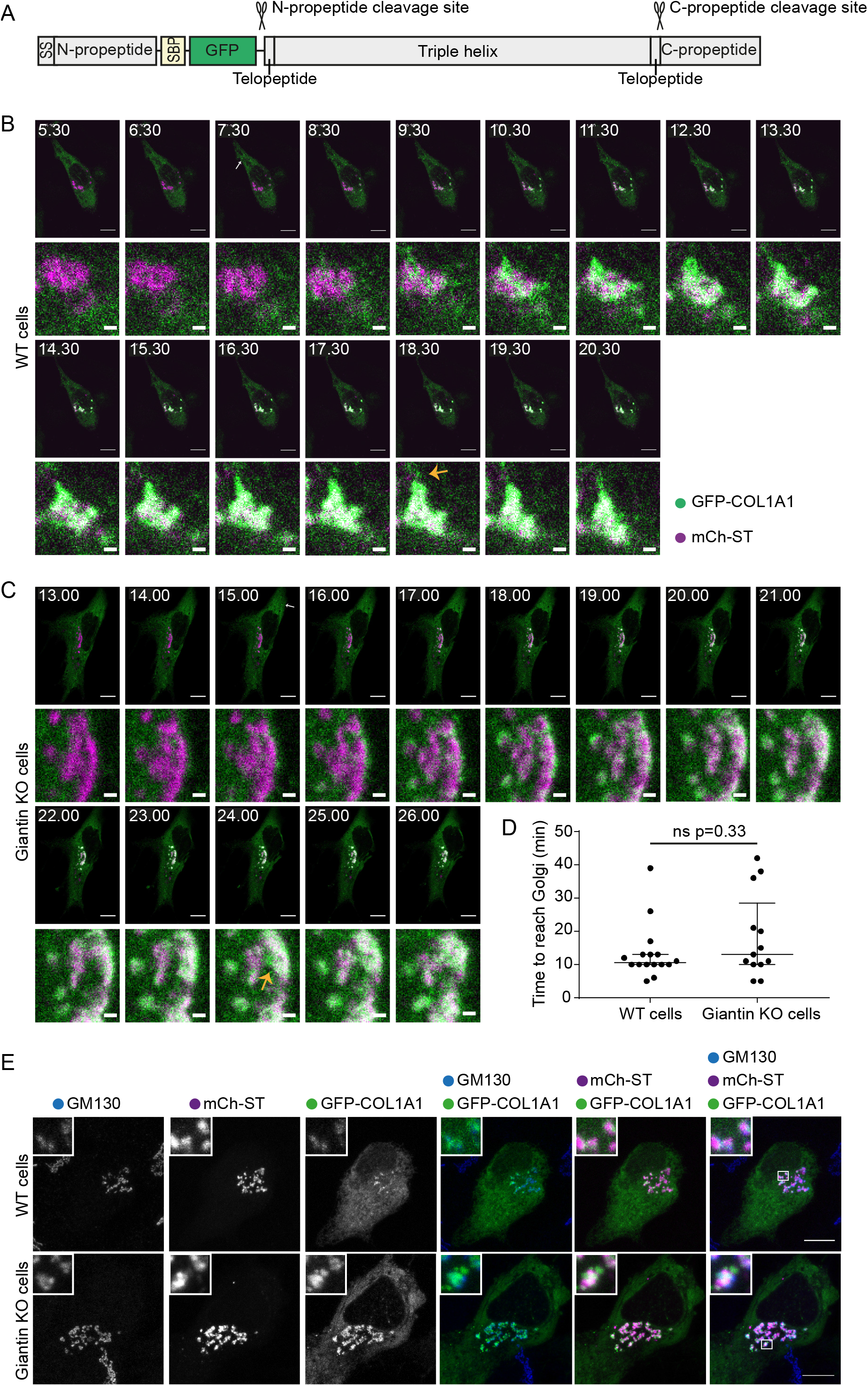
ER-Golgi trafficking of pro-α1(I) is unperturbed in giantin KO cells. **A** Schematic of the pro-SBP-GFP-COL1A1 construct used in this study. SS = signal peptide SBP = streptavidin binding protein. **B-C**. Single plane confocal stills taken from live imaging movies of a (A) WT and (B) giantin KO cell stably expressing GFP-COL1A1 (green) and transiently transfected with mCh-ST (magenta) and an ER RUSH hook. Frame times in the top left corner indicate the time (minutes) post biotin addition which releases GFP-COL1A1 from the ER hook. Scale bars 10 μm on overview and 1 μm on zoom. White arrows indicate the appearance of peripheral GFP-COL1A1 punctate. Orange arrows indicate emerging post-Golgi carriers. See also supplemental movies. **D.** Quantification of the time taken for GFP-COL1A1 to first appear at the Golgi in movies represented in (A-B). Each dot represents one cell (16 WT cells and 13 giantin KO cells) and movies were collected over 7 independent experiments. Bars represent median and interquartile range. Results are not significant using a Mann-Whitney-U test. **E.** Maximum projections of confocal z stacks of WT and giantin KO cells imaged live as in (A-B) and then fixed with PFA as soon as the GFP-COL1A1 appeared around the Golgi. Cells were then immunolabelled for the Golgi marker GM130 (blue). Scale bars 10 μm.

In WT cells, the GFP-COL1A1 first accumulates around the mCh-ST positive Golgi compartment upon release (Figure 3B, Supplemental Movie 1). Fixation at this timepoint followed by immunolabelling of Golgi markers confirmed that this predominantly represents the GFP-COL1A1 entering the *cis*-Golgi (Figure 3E). Shortly after *cis*-Golgi filling, the GFP-COL1A1 progresses through the Golgi into the mCh-ST-positive cisternae before leaving the Golgi in tubular vesicular carriers (Figure 3B-C arrow). This “short loop” pathway from the ER to cis-Golgi in the absence of discernible carriers has been described previously (McCaughey et al., 2019). Crucially here, we found that GFP-collagen also traverses this pathway in the giantin KO cells (Figure 3C, Supplemental movie 2).

To measure the kinetics of ER-to-*cis*-Golgi transport we quantified the time between biotin addition and the appearance of the GFP-COL1A1 in the *cis*-Golgi. Trafficking rates were equivalent in both cell lines (Figure 3D). We therefore conclude that the collagen phenotypes seen in giantin KO cells are not due to the use of an alternative trafficking route or faster kinetics.

### Giantin KO cells exhibit defects in processing of type I procollagen

During these experiments we observed that giantin KO cells stably expressing GFP-COL1A1 were often surrounded by extracellular GFP-positive fibres. Immunofluorescence with a COL1A1 antibody identified these as collagen fibrils, indicating that the GFP-tag is incorporated into the ECM (Figure 4A). No GFP-COL1A1 positive fibrils could be detected in the matrix of WT cells despite an abundance of collagen and the expression of GFP alone did not label the ECM.

**Figure 4.**
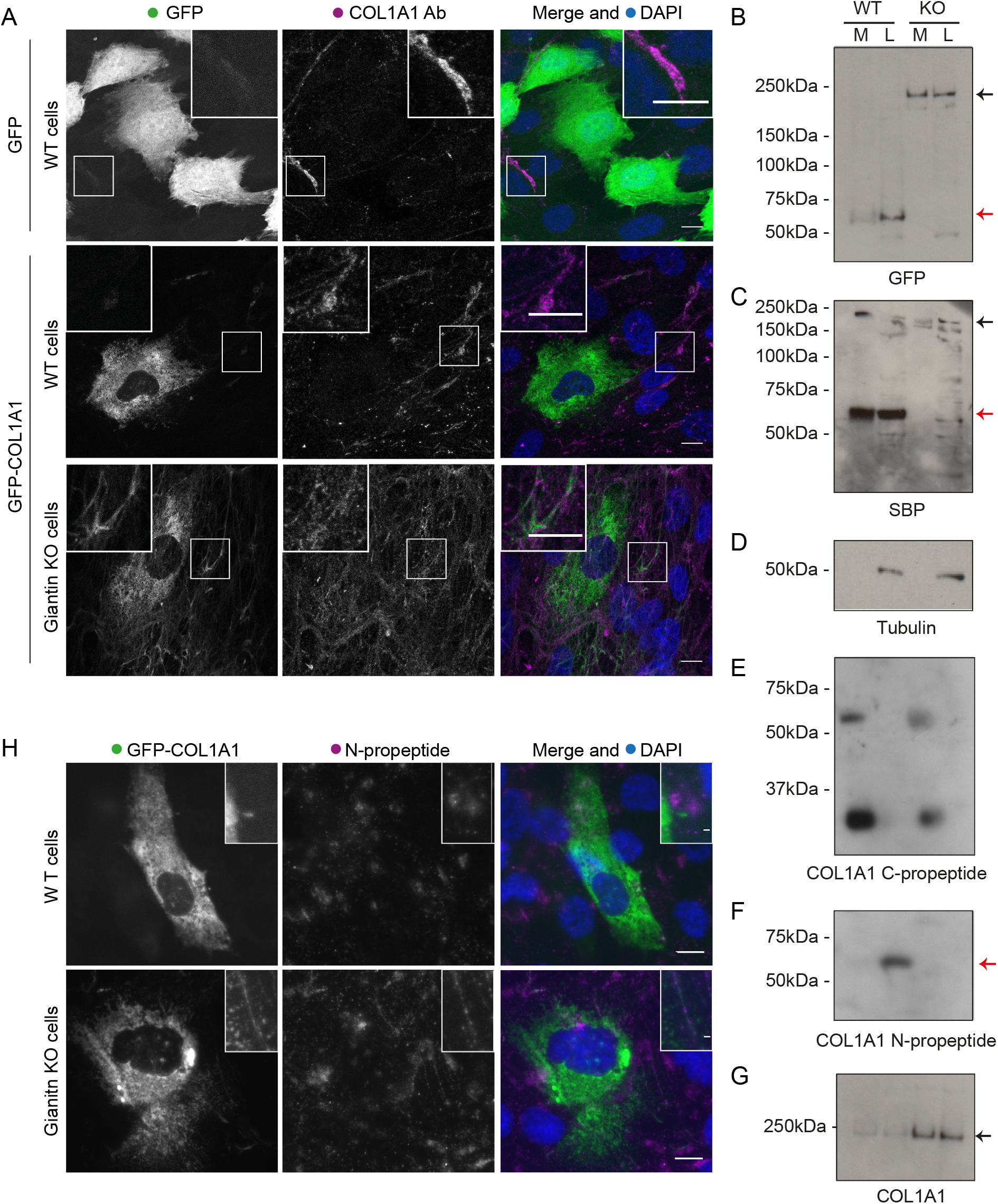
Procollagen processing is defective in giantin KO cells. **A.** Representative images of WT and giantin KO cell expressing GFP-COL1A1 or GFP alone as indicated and immunolabelled for pro-α1(I) (COL1A1, magenta). Nuclei are stained with DAPI (blue). Images are maximum intensity projections of widefield z-stacks. Scale bars 10 μm. **B-G.** Western blots of secretion assays showing the medium (M) and lysate (L) fractions of WT and giantin KO cell cultures immunoblotted for (B) GFP, (C) SBP, (D) Tubulin, (E) C-propeptide (LF41 antibody) (F) N-propeptide (LF39 antibody) and (G) pro-α1(I) (COL1A1). Black arrows indicate full length procollagen and red arrows highlight N-propeptide bands. **H.** Maximum intensity projection widefield z-stacks of WT and giantin KO cells stably expressing GFP-COL1A1 immunolabelled for the type I procollagen N-propeptide (LF39 antibody). Scale bars 10 μm.

In our reporter construct, GFP is inserted upstream of the N-propeptide cleavage site and should be removed during processing (Figure 3A). The presence of GFP in the ECM therefore implies that either N-propeptide processing is defective or that free N-propeptides are abnormally interacting with collagen fibrils. To investigate this, we assayed secretion from these stable cell lines by immunoblotting. In both the media and lysate fractions of WT cultures, a dominant GFP-positive band was discernible at ~60kDa, which corresponds to the size of the tagged N-propeptide (Figure 4B). In KO cells, however, the dominant GFP-positive band was detected at ~200kDa. This is consistent with the size of unprocessed, tagged procollagen. No band was discernible at ~60kDa in the KO cells, even after enrichment by immunoprecipitation (Supplemental Figure 2A) suggesting N-propeptide cleavage does not take place. These results were supported by additional immunoblots detecting the SBP tag (Figure 4C). In this instance, full length procollagen is also evident in WT cells, likely due to the higher affinity of the SBP antibody, but the N-propeptide remains undetectable in the KO cells.

To verify our interpretation of these observations with respect to the N-propeptide, we performed immunoblots using antibodies specifically targeting the N- and C-propeptide domains of pro-α1(I). C-propeptide cleavage was comparable in WT and KO cells (Figure 4E) with bands of ~60 kDa and ~30 kDa detected. However, an antibody against the N-propeptide detected a band of ~60kDa in WT cell lysates that was again completely absent in the KO cultures (Figure 4F). This N-propeptide band was also detected in the media of WT cells but only following enrichment by immunoprecipitation (Supplemental Figure 2A). It is therefore present but in low quantities. Full length collagen protein levels were again higher in the KO cells (Figure 4G).

We also used the N-propeptide-specific antibody (LF-39) to immunolabel the ECM of WT and KO cells. Whilst this antibody failed to label the CDM of WT cells, it did label fibrils in the KO cell matrix where it colocalised with the extracellular GFP signal (Figure 4H). Together these data indicate that the presence of GFP in the collagen matrix of KO cells is due to a failure in N-propeptide cleavage. The above observations are also consistent with N-propeptide cleavage taking place intracellularly in WT cells.

Secretion of unprocessed procollagen has been reported in the absence of Hsp47 (Ishida et al., 2006), however, Hsp47 levels in the KO cells were normal, ruling this out as a cause (Supplemental Figure 2B-C). We also hypothesised that the absence of N-propeptide in the KO cells could be due to its increased degradation, but treatment of KO cells with proteasomal and lysosomal inhibitors did not render it detectable (Supplemental Figure 2D). Interestingly, N-propeptide levels in the WT cells did change upon treatment with these inhibitors, thus confirming the presence of an intracellular pool sensitive to these pathways (Supplemental Figure 2D).

### N-terminal processing of type I procollagen is defective in other giantin KO lines

Since the above data were derived from a single clone of *GOLGB1* KO cells, we generated new KO cell lines in which to validate our findings. This time we targeted exon 13 of the *GOLGB1* gene (instead of exon 7) and used the lentiCRISPRv2 system (Sanjana et al., 2014; Shalem et al., 2014) for transfection in order to vary our approach. We obtained three clones with different indel mutations (Supplemental Figure 3). Immunoblots with a giantin polyclonal antibody confirmed the loss of full-length protein, although hints of a potential truncated form could be seen in low abundance (Figure 5A). Immunofluorescence staining with antibodies targeting the N- and C-termini of giantin was unable to detect any protein (Figure 5B). As previously described for the original knockout line (Stevenson et al., 2017), no gross changes in Golgi morphology or ER exit site abundance were apparent following loss of giantin (Figure 5B).

**Figure 5.**
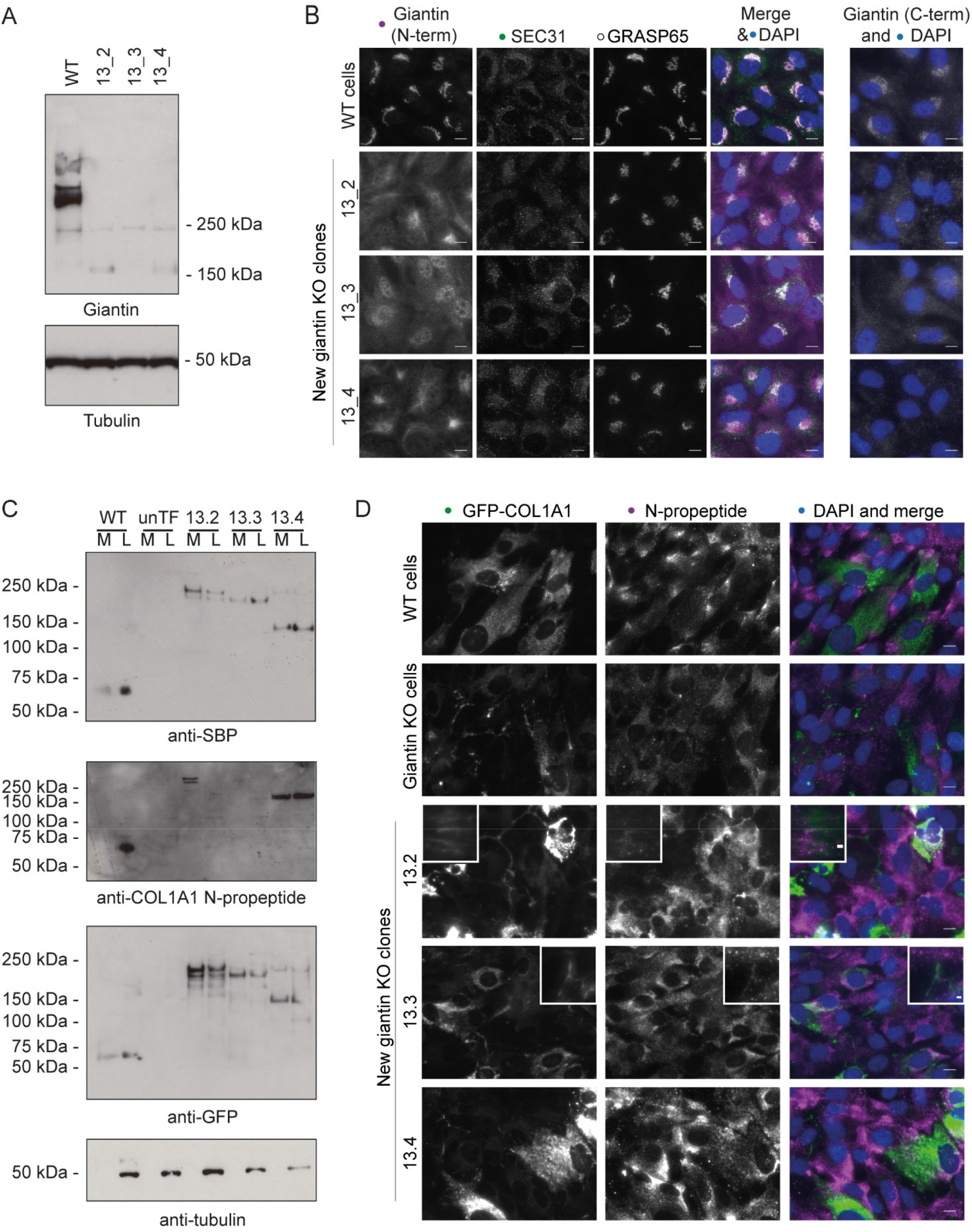
Validation of processing defects in other giantin mutant lines. **A.** Western blot of WT RPE1 and giantin exon 13 KO clone lysates probed for giantin (polyclonal against full length protein) and tubulin (housekeeping) **B.** WT and giantin KO RPE1 cell clones immunolabelled for the N- and C-terminus of giantin, ER exit sites (Sec31) and the Golgi (GRASP65) as indicated. Scale bar 10 μm. **C.** Western blots of secretion assays performed on WT and giantin KO clones stably expressing GFP-COL1A1 and untransfected giantin KO cells (unTF). Blots show the medium (M) and lysate (L) fractions of each cell population immunoblotted for GFP, SBP and type I procollagen N-propeptide as indicated. **D**. Maximum projection widefield images of GFP signal in giantin KO clones stably expressing GFP-COL1a1 and immunolabelled for pro-α1(I) N-propeptide (magenta). Scale bar 10 μm.

Using the exon 13 mutant clones we derived new stable cell lines expressing GFP-COL1A1. We also made a new WT GFP-COL1A1 stable line to be sure of the WT phenotype. As with our original clone, we were not able to detect cleaved N-propeptide in either the media or lysate fractions from these new KO cell cultures (Figure 5C). We did however note some variability in the molecular weight of the procollagen bands detected between clones, perhaps suggesting additional processing defects or deficiencies in post-translational modifications. Imaging of the new KO clones again also confirmed incorporation of the GFP-tag and the N-propeptide into the extracellular collagen matrix in two of these lines (Figure 5D). The phenotypes of the new WT stable cell lines also validated those of the original. Overall, these results confirm that N-propeptide processing is defective in the absence of giantin.

## Discussion

In this study we demonstrate for the first time that giantin is required for N-terminal, but not C-terminal, processing of type I procollagen. In its absence, procollagen intermediates still bearing the N-propeptide are secreted and incorporated into the ECM. Unlike the C-propeptide, retention of the N-propeptide does not preclude fibril formation (Hulmes et al., 1989; Miyahara et al., 1984; Miyahara et al., 1982; Romanic et al., 1992) and thus it is not surprising that a collagen fibrils are formed. We also found that free N-propeptide is clearly detected in the cell layer of WT cells, implying it is cleaved inside the cell. Altogether, these data indicate the presence of a giantin-dependent intracellular pathway for procollagen processing.

The most likely cause of the observed processing deficiency in KO cells is a defect in the expression, trafficking, or processing of an N-proteinase. Although we have not yet identified any trafficking defects in our giantin KO cells either generally (Stevenson et al., 2017) or specifically for procollagen, it is possible that other cargoes such as processing enzymes will be affected. To date five enzymes have been implicated in the N-terminal processing of type I procollagen - ADAMTS2, ADAMTS3, ADAMTS14 (Bekhouche and Colige, 2015; Colige et al., 1997; Colige et al., 2002; Fernandes et al., 2001), meprin-α and meprin-β (Broder et al., 2013). Our previous RNAseq analysis of RPE1 cells (ArrayExpress E-MTAB-5618) failed to detect any ADAMTS2 transcript, even in WT cells, however ADAMTS3 and −14 mRNA was present and interestingly their expression was down-regulated following giantin KO (Stevenson et al., 2017). Unfortunately, our attempts to further this line of enquiry have so far been unfruitful due to a lack of suitable reagents.

A second possible cause of defective processing is glycosylation deficiency. Indeed, it has been shown that C-proteinase activity is sensitive to inhibitors of *N-*glycosylation (Duksin et al., 1978). Loss of giantin function affects the enzyme composition of the Golgi and the *O-*glycosylation of specific substrates (Lan et al., 2016; Stevenson et al., 2017). It is thus feasible that at least one of the N-propeptidases is incorrectly glycosylated in giantin KO cells in such a way as to impede activity. Alternatively, procollagen itself may be incorrectly glycosylated preventing its identification as a substrate. We did perform a global analysis of glycoproteins by mass spectrometry but, as predicted from previous results (Stevenson et al., 2017), there were no differences in the major glycosyl chains produced by WT or KO cell lines. This approach did not permit us to determine the glycosylation of specific substrates.

A crucial observation made in this study is that N-propeptide processing occurs, at least in part, inside the cell since cleaved N-propeptide, but not C-propeptide, is readily detectable in the lysates of WT cells. Whilst it is possible that this pool of N-propeptide is in fact extracellular N-propeptide associated with the cell surface or ECM, we rule this out for several reasons. Firstly, care was taken to lyse the cells gently and leave the ECM behind. Secondly, N-propeptide in the cell layer fraction was sensitive to inhibitors of both lysosomal and proteasomal degradation. Thirdly, N-propeptides have been noted to have biological functions consistent with intracellular localisation (Oganesian et al., 2006). Finally, there is precedent for the detection of intracellular procollagen processing in both chick and mouse tendons (Canty-Laird et al., 2012; Canty et al., 2004; Humphries et al., 2008). In these studies, processed forms of type I procollagen were detected in a detergent soluble fraction from tendon explants, consistent with an intracellular pool. Furthermore, inhibition of procollagen secretion by BFA treatment, which causes the cis- and medial-Golgi machinery to accumulate in the ER, does not prevent N-terminal processing, indicating procollagen can be processed in the early secretory pathway where giantin is located (Canty-Laird et al., 2012). Considering this, together with the data presented here, we conclude that giantin is a crucial component of an intracellular pathway of procollagen processing.

Classically, human patients with mutations affecting N-terminal processing of type I procollagen suffer from Ehlers-Danlos syndrome (EDS) type VII, a disease characterised by joint hypermobility, skin hyperextensibility, and dislocations (Byers et al., 1997; Colige et al., 1999; Giunta et al., 1999; Smith et al., 1992). Collagen fibres in these patients tend to be thinner and have irregular contours (Malfait et al., 2013; Van Damme et al., 2016). ADAMTS2 deficient mice also have thinner collagen fibrils in the skin (Li et al., 2001). This is not what we observe in the CDM of *GOLGB1* KO cells and the characteristics of EDS patients have not been described in *GOLGB1* KO animals or humans. There are several reasons why giantin loss may not mimic an EDS phenotype. Firstly, prior RNAseq data shows that the expression of many matrix components is altered following loss of giantin (Stevenson et al., 2017). We believe this may be compensatory in both cells and zebrafish and must affect matrix quality more globally. Giantin also impacts ciliary function which would compound phenotypes (Asante et al., 2013; Stevenson et al., 2017). Secondly, we do not exclude the possibility that N-terminal processing still occurs in the extracellular space in our systems. In fact, we think it likely given the mild impact on matrix organisation observed. In cases of EDS type VII, mutations in ADAMTS2 or the N-terminus of pro-α1(I) directly impact N-propeptide cleavage regardless of location or environmental conditions. We propose giantin acts more indirectly to regulate an intracellular processing pathway which acts in concert with other complimentary mechanisms, such as extracellular cleavage, for co-ordinated and efficient processing under different conditions. This raises the intriguing possibility of specific pools of collagen that require intracellular processing for specific functions, such as fracture repair.

We found that *COL1A1* gene expression is reduced in both *golgb1^X3078/X3078^* mutant zebrafish and KO cells (Stevenson et al., 2017). This contrasts with protein levels which are more abundant *in vitro*, at least in the cells we have analysed here. Previous studies have suggested that the pro-α1(I) N-propeptide is capable of acting as a negative regulator of pro-α1(I) synthesis (Horlein et al., 1981; Oganesian et al., 2006; Paglia et al., 1979; Wiestner et al., 1979). Importantly, the point of inhibition occurs during translational chain elongation or termination (Horlein et al., 1981; Oganesian et al., 2006; Paglia et al., 1979). Thus, we propose that despite the reduced abundance of mRNA transcript, translation of pro-α1(I) is more efficient in giantin KO cells in the absence of inhibitory N-propeptide. Consistent with reports that only endogenously expressed N-propeptide can inhibit translation, we were unable to rescue our phenotypes by supplementing media with recombinant type I procollagen N-propeptide.

As previously described, *golgb1^X3078/X3078^* mutant zebrafish show developmental growth delay, elongated craniofacial structures and ectopic mineralisation of soft tissue (Stevenson et al., 2017). Here, we additionally report that mutant adult caudal fins accumulate significant numbers of fractures and show defects in the maturation of newly deposited matrix during fracture repair. Our observations of increased calcein labelling in mutant fish fractures is consistent with either increased osteoid formation or hyper-mineralisation. We have previously reported that expression of the glycosyltransferase GALNT3 is reduced in both *GOLGB1* KO cells and zebrafish (Stevenson et al., 2017). GALNT3 is required to glycosylate and activate FGF23, which is an inhibitor of bone mineralisation (Wang et al., 2008). The observed defects in fracture repair could therefore be the result of FGF23 deficiency.

Alternatively, or perhaps additionally, collagen N-propeptides have been linked to the differentiation and apoptosis of osteoblasts (Oganesian et al., 2006) and osteoclasts (Hayashi et al., 2011) respectively. We have not been able to directly demonstrate that the procollagen processing defects observed in cells translate to fish, however it is interesting to speculate that processing defects may affect the balance of bone synthesis and resorption in the mutant fractures. The significant impact of the loss of giantin on mineralisation likely explains why mutant animals primarily present with skeletal defects and other type I collagen rich tissues, such as skin, are less affected.

The observed increased incidence of fractures in mutants could result from an increased frequency of fractures in brittle bones or an accumulation of fractures due to slow healing. Although fracture repair is delayed in the mutants, our data show that the described differences between WT and mutant fractures largely resolve within three weeks. This seems insufficient to account for number of spontaneous fractures that accumulate, given that callouses can persist for months overall (Geurtzen et al., 2014). Combined with the clustered nature of a significant number of fractures we believe it likely that the bones are more brittle. This would be consistent with a defect in type I collagen deposition, as mutations in the *COL1A1* gene are associated with osteogenesis imperfecta in both humans (Glorieux, 2008) and zebrafish (Fisher et al., 2003; Gistelinck et al., 2018). Some N-propeptide processing deficiencies can cause disease with characteristics of both osteogenesis imperfecta and EDS in humans (Cabral et al., 2005; Makareeva et al., 2006). Giantin KO also affects the abundance of other matrix proteins affecting bone quality (Stevenson et al., 2017). Further study will be needed to define which, if any, phenotypes seen in zebrafish are a direct result of defective N-terminal processing of type I procollagen.

## Acknowledgements

We would like to thank Franck Perez and Gaelle Boncompain (Institut Curie, France) for sharing the RUSH system with us, Helen Dawe (University of Exeter, UK) and Stuart Haslam (Imperial College London, UK) for their technical help and advice, Andrew Herman and the UoB flow cytometry facility for sorting our cell lines, Kate Heesom and the UoB proteomics facility for performing exploratory mass spec runs, Lucy McGowan for her assistance with the fish experiments, and Alain Colige (University of Liege, Belgium) for his help and advice and for providing an ADAMTS2 construct. This work was funded by the MRC (MR/P000177/1) and Versus Arthritis (21937 and 22044). We also thank the MRC and Wolfson Foundation for establishing the Wolfson Bioimaging Facility, and confocal microscopy was supported by a BBSRC ALERT 13 capital grant (BB/L014181/1). Thanks to Karl Kadler for critical reading of the manuscript. The authors declare no competing financial interests.

## Author Contributions

Conceptualisation D.J.S & N.L.S. Formal analysis N.L.S. Funding acquisition and supervision D.J.S & C.L.H. Investigation and methodology N.L.S and D.J.M.B. Project administration D.J.S. Visualisation N.L.S. Writing original draft N.L.S and D.J.S. Writing review and editing NLS, D.J.M.B, D.J.S, and C.L.H.

## Supplemental Figures

**Supplemental Figure 1.**
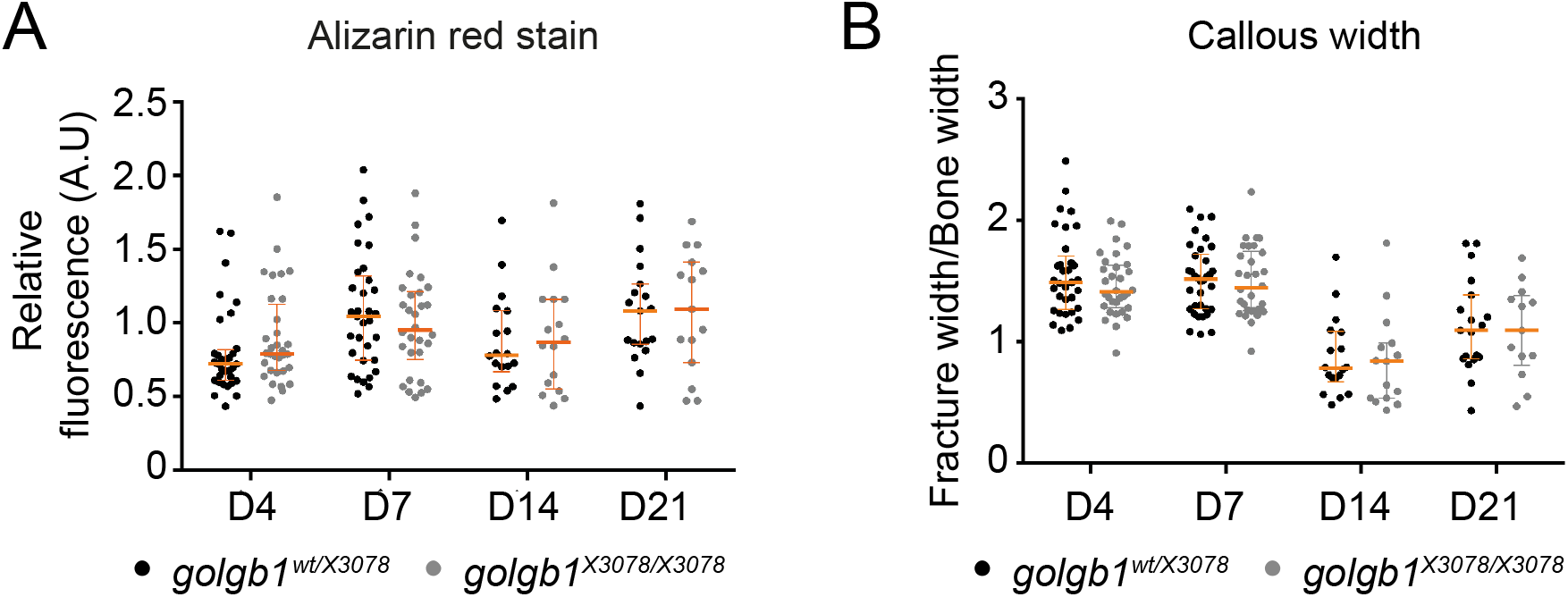
Fracture analysis in WT and mutant zebrafish. **A-B.** Quantification of (A) the intensity of alizarin red staining and (B) callous width of experimentally induced fractures normalised to adjacent healthy bone at different timepoints post injury (D=days post injury) in *golgb1^wt/X3078^* and *golgb1^X3078/X3078^* fish. Dots represent individual fractures. At timepoints 4 and 7 d.p.f eleven fish per line were quantified. At timepoints 14 and 21 d.p.f n= 6 HETs and n=4 HOM fish. All from a single experiment. Horizontal and vertical orange lines depict median and interquartile range respectively. P value calculated with a Mann Whitney test comparing the mean value per fish (three fractures per fish).

**Supplemental Figure 2.**
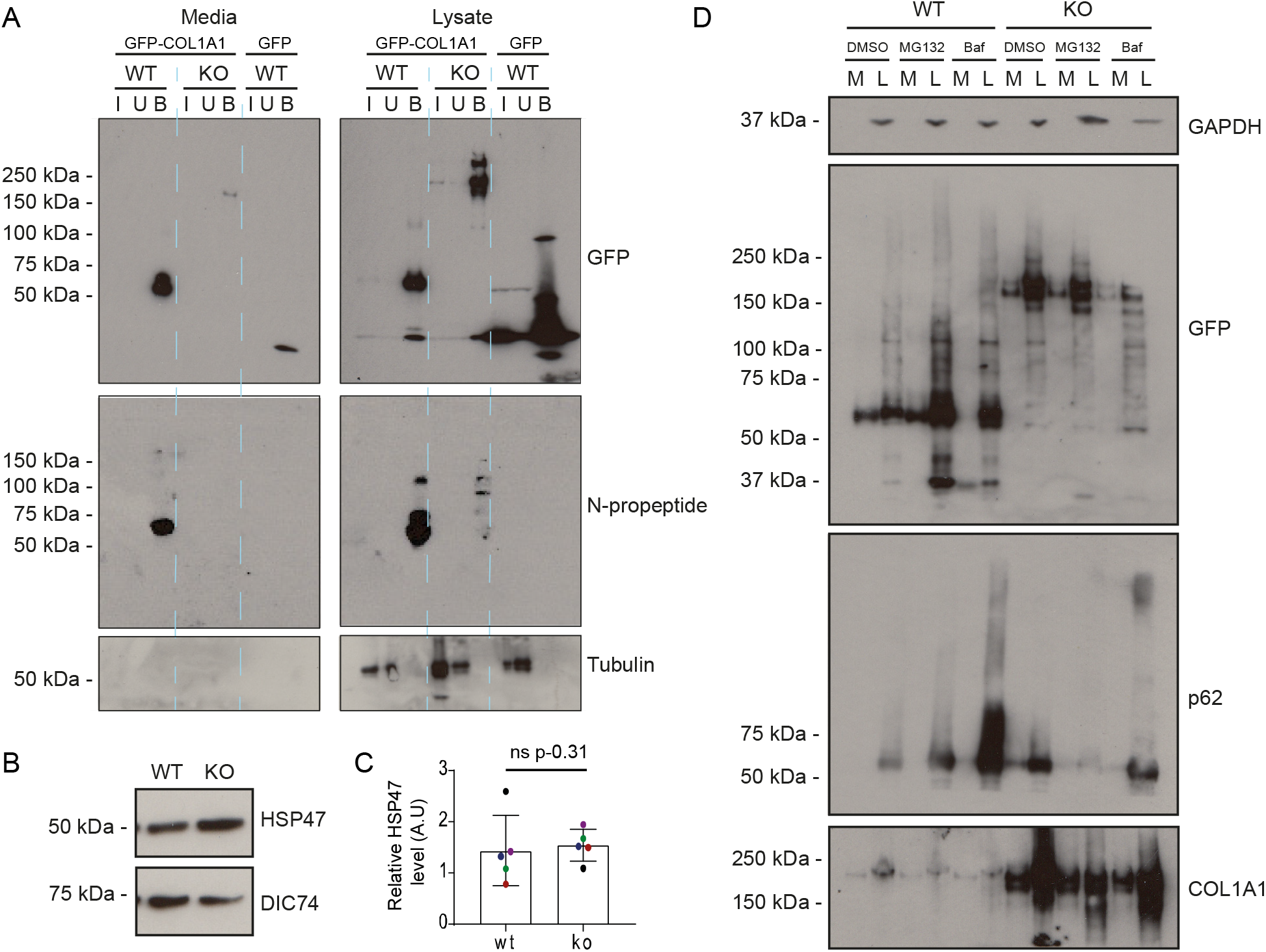
Procollagen processing controls. **A**. Western blot of a GFP trap of media (M) and lysates (L) fractions of WT and KO cell cultures. Cells were either expressing GFP-COL1A1 or GFP alone as indicated. Blots show the input (I), unbound (U) and bound (B) fractions of the IP immunoblotted for GFP, type I procollagen N-propeptide (LF39 antibody), and tubulin (housekeeping). **B**. Western blot of HSP47 and DIC74 (housekeeping) in WT and giantin KO RPE1 cell lysates. **C.** Densitometry of the semi-quantitative ECL western blots represented in (A). HSP47 levels are normalised to DIC74. Each dot represents an independent biological replicate and replicates are colour coded between cell lines. Bars show median and interquartile range (n=5 biological replicates). P value calculated with a Mann Whitney test. **D.** Western blots of media (M) and lysate (L) fractions of WT and giantin KO cell cultures stably expressing GFP-COL1A1 following treatment with DMSO (vehicle control), MG132 or bafilomycin (baf). Blots are probed for GAPDH (housekeeping), GFP (GFP-COL1A1), p62 (positive control for bafilomycin) and pro-α1(I) (COL1A1) as indicated.

**Supplemental figure 3.**
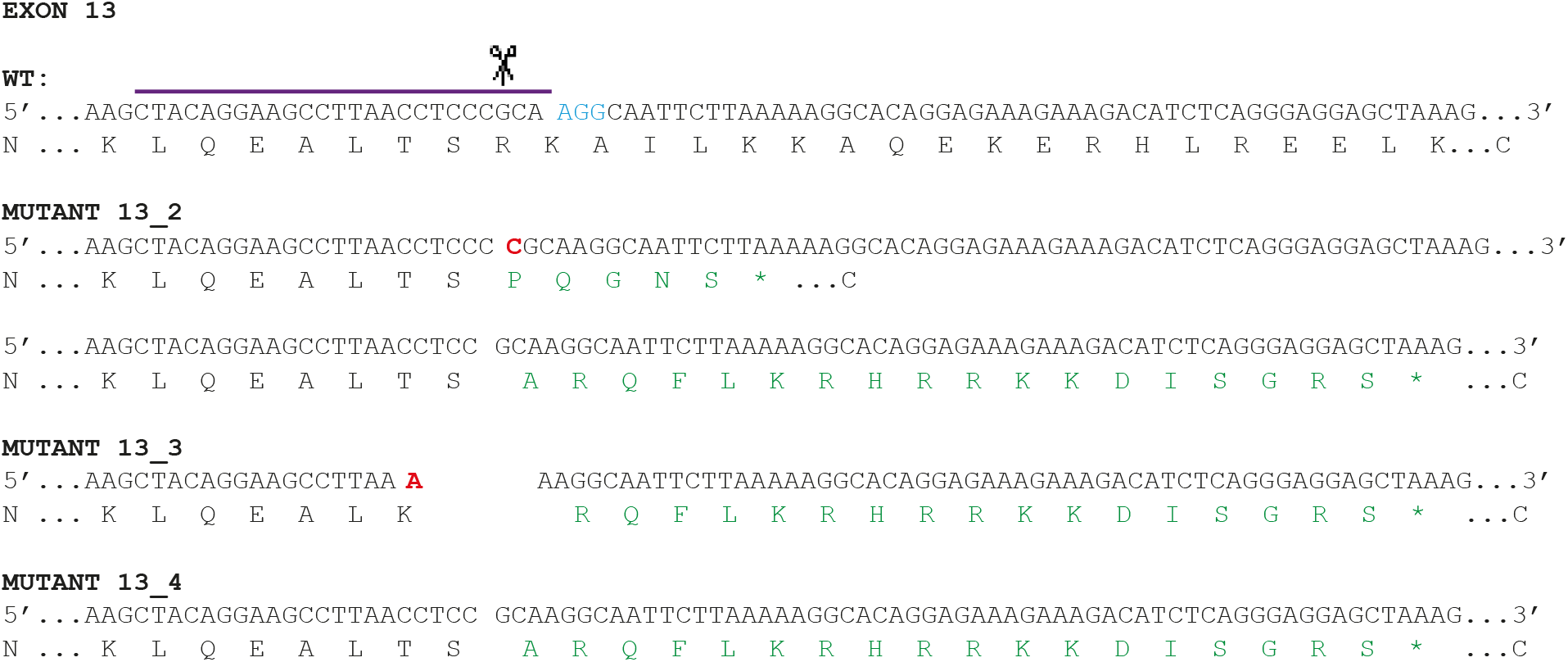
Validating pro-α1(I) processing defect in other giantin KO clones. Genomic DNA sequence and amino acid translation at CRISPR/Cas9 mutation site in three new giantin KO clones targeted at exon 13. On the WT sequence the gRNA sequence is indicated with a purple line, the cut site is indicated by scissors and PAM site is in blue text. In mutant sequences red text indicated inserted base pairs in genomic sequence and green text shows altered amino acids. Asterisks denote premature stop codons.

## Legends to Supplemental Movies

**Supplemental Movie 1**

Representative movie of a RUSH assay in a WT RPE1 cell stably stably expressing GFP-COL1A1 (green) and transiently transfected with mCh-ST (magenta, Golgi label) and an ER RUSH hook. At time 0 biotin and ascorbic acid were added to the medium to release the collagen from the hook and promote procollagen folding respectively to trigger anterograde trafficking. Movie is cropped to the relevant time frame – time is indicated in top left corner as mm:ss. Frames were captures as single frame images every 20 seconds. Movie frame rate 12 frames/s. Scale bar 10 μm.

**Supplemental Movie 2**

Representative movie of a RUSH assay in a Giantin KO RPE1 cell stably expressing GFP-COL1A1 (green) and transiently transfected with mCh-ST (magenta, Golgi label) and an ER RUSH hook. At time 0 biotin and ascorbic acid were added to the medium to release the collagen from the hook and promote procollagen folding respectively to trigger anterograde trafficking. Movie is cropped to the relevant time frame – time is indicated in top left corner as mm:ss. Frames were captures as single frame images every 20 seconds. Movie frame rate 12 frames/s. Scale bar 10 μm.

## Methods

### Zebrafish husbandry and genetic lines

London AB zebrafish were maintained using standard conditions (Alestrom et al., 2019). Ethical approval was obtained (University of Bristol Ethical Review Committee) and experiments performed under Home Office Project License number 30/3804. The *golgb1^bsl077^* (henceforth *golgb1^X3078^*) allele is described in (Stevenson et al., 2017). The *golgb1^X3078^* allele was crossed with fish carrying the *Tg(col1a1a:EGFP)^zf195tg^* (henceforth *col1a1a:GFP*) transgenic reporter of the 1.4 kb *col1a1a* promoter region (Kague et al., 2012). The GFP positive *golgb1^wt/X3078^* + *col1a1a:GFP* individuals were in-crossed to generate homozygotes. In all experiments, homozygotes were compared to siblings.

### Fin fracture assays

Fin fractures were performed as previously described (Geurtzen et al., 2014). Briefly, 7-month-old adult fish were anaesthetised with MS222 (Sigma-Aldrich) and moved into a petri dish with the head placed on a bed of tissue soaked in anaesthetic and the tail splayed on the dish for imaging. Caudal fins were imaged prior to injury and any pre-existing fractures counted. Fractures were induced by pressing on an individual segment of bone in the tail rays with the end of a semi-flexible plastic pipette tip. Four fractures per fish were introduced in non-pigmented sections of fin in the distal third of the tail, avoiding the four most dorsal and ventral rays. Fish were then reimaged live at various time points post-injury.

### Live staining of bone

To visualise bone repair in live zebrafish, Alizarin Red stain (ARS) and calcein green stain was used. ARS staining solution contained 74 μM Alizarin Powder (Sigma-Aldrich) and 5 mM HEPES dissolved in Danieau’s buffer. Calcein green staining solution contained 40 μM calcein powder (Sigma-Aldrich) dissolved in Danieau’s buffer and was pH calibrated to pH 7.4. Live fish were immersed in either ARS or calcein green for two hours, then immersed in fresh system water for 15 minutes prior to imaging to clear excess stain from the fish.

Fluorescence intensity at fracture sites was measured using image j – lines were drawn along the fracture callous length and then the line width expanded to encompass the whole fracture. The mean intensity was then measured across this area, before using the same ROI to measure the mean intensity of adjacent, uninjured hemi-ray. Fracture intensity was then normalised against the healthy bone to account for differences in baseline expression caused by reporter integration number.

### Cell lines and culture

Human telomerase-immortalised retinal pigment epithelial cells (hTERT-RPE1, ATCC) were grown in DMEM-F12 (Life Technologies, Paisley, UK) supplemented with 10% decomplemented FCS (Gibco). Cell lines were not authenticated after purchase other than confirming the absence of mycoplasma contamination. The main *GOLGB1* KO hTERT-RPE1 cell line used is described in (Stevenson et al., 2017).

New *GOLGB1* KO cell lines with mutations in exon 13 were generated using the lentiCRISPRv2 system (lentiCRISPR v2 was a gift from Feng Zhang (Addgene plasmid # 52961; http://n2t.net/addgene:52961; RRID: Addgene 52961)). Guide RNA sequences were designed using Benchling (http://benchling.com) and inserted into the lentiCRISPRv2 construct as described in (Sanjana et al., 2014; Shalem et al., 2014) to be co-expressed with Cas9. [Target sequences used: 5’ - CCACCGGGAAGCCTTAACCTCCCGCA - 3’ and 5’ -- AAACTGCGGGAGGTTAAGGCTTCCC – 3’]. The cloned plasmid was packaged into lentivirus using lenti-X-packaging-single-shots (VSVG) (Takara, #631275) according to the manufacturer’s instructions. After harvest, 1 ml virus supernatant was added to 80% confluent RPE1 cells in a 6cm dish (after removal of growth media) in the presence of 8 μg/ml polybrene. Virus was incubated for 1 h before adding back fresh growth medium supplemented with 8 μg/ml polybrene. Transfection medium was replaced after 24 h. Cells were passaged 24 h later in 20 μg/ml puromycin to select transfected cells. After seven days, individual cells were FACS sorted into a 96 well plate to grow up clones and screen for *GOLGB1* knockout by immunofluorescence and western blot. To identify the mutations, genomic DNA was extracted from each clone using the Purelink genomic DNA mini kit (Invitrogen) and the region targeted by the gRNAs was amplified by PCR [primers: forward 5’ - GCTGGCAGCTGAAGAGCAATTCCA - 3’ reverse 5’ - GTTGAGTGTGATGCTGTTCTGTGGCT - 3’]. PCR products were cloned into the pGEM-T Easy vector according to the manufacturer’s instructions (Promega) and sequenced using predesigned primers against the T7 promoter (MWG Eurofins).

Stable cell lines were made by lentiviral transduction as described above using the Lenti-X Single shot lentivirus packaging system (Takara, #631275) in combination with the lentiviral vector pLVXpuro-proSBP-GFP-COL1A1 (Addgene #110726) described in (McCaughey et al., 2019). To ensure only low levels of over-expression all cell lines were FACs sorted. GFP only stable cells are described in (Asante et al., 2014).

Transient transfections were performed using Lipofectamine 2000 according to the manufacturer’s instructions using 2 μg DNA for a 35 mm well (Invitrogen, Carlsbad, CA, Cat# 11668027). Plasmid: Str-KDEL-IRES-ST-mCherry is described in (McCaughey et al., 2019) (Addgene; no. 110727).

### Antibodies

Rabbit anti-COL1A1 (Novus Cat# NB600-408, RRID:AB_10000511), mouse anti-DIC74 (Millipore Cat# MAB1618, RRID:AB_2246059), mouse anti-HSP47 (Enzo Life Sciences Cat# ADI-SPA-470, RRID:AB_10618557), mouse anti-GM130 (BD Biosciences Cat# 610823, RRID:AB_398142), rabbit anti-giantin (N-terminus, BioLegend Cat# 924302, RRID:AB_2565451), rabbit anti-giantin (C-terminus, gift from Martin Lowe), mouse anti-SBP clone 20 (Millipore Cat# MAB10764, RRID:AB_10631872), rabbit polyclonal anti-N- and C-propeptide of COL1A1 (LF-39 and LF-41 respectively, both gifts from Larry Fisher, National Institutes of Health, Bethesda, MD;(Fisher et al., 1995)), mouse monoclonal anti-GFP (Covance Cat# MMS-118P-500, RRID:AB_291290), sheep polyclonal anti-GRASP65 (gift from Jon Lane), tubulin (Sigma-Aldrich Cat# T5168, RRID:AB_477579), mouse monoclonal anti-Sec31A (BD Biosciences Cat# 612350, RRID:AB_399716), mouse p62 (Novus Cat# H00008878-D01P, RRID:AB_1504204). Secondary HRP-conjugated antibodies were obtained from Jackson ImmunoResearch. Alexa-Fluor^®^ conjugated secondaries were obtained from Invitrogen.

### Immunofluorescence

Cells were grown on autoclaved coverslips (0.17 mm thickness #1.5 Thermo Fisher Scientific) prior to fixation with 4% PFA/PBS for 10 minutes, permeablilsation with 1% Triton X-100/PBS for 10 minutes and blocking with 3% BSA/PBS for 30 minutes. Sequential incubations with primary and secondary antibodies diluted in block were performed for one hour, washing in PBS in between. Nuclei were labelled with DAPI (4,6-diamidino-2-phenylindole; Life Technologies, D1306) for 3 minutes. Coverslips were mounted in MOWIOL 4-88 (Merck-Millipore). All steps were conducted at room temperature and in the dark post-secondary antibody addition. When labelling for extracellular matrix, the triton permeability step was excluded from the protocol.

For widefield microscopy an Olympus IX-71 inverted microscope was used with a 60×1.42 NA oil-immersion lens, Exfo Excite xenon lamp illumination, and single-pass excitation, emission and multipass dichroic (Semrock) filters. Images were captured on an Orca-ER CCD (Hamamatsu). The system was controlled using Volocity (v.5.4.1; PerkinElmer). Chromatic shifts in images were registration corrected using TetraSpek fluorescent beads (Thermo Fisher Scientific). Fixed cell imaging by confocal microscopy was performed using a Leica SP5II system at 1024×1024 xy resolution. Tilescans were performed to capture larger areas of the CDM. On both systems, image stacks were taken with Δz of 0.2 μm and, unless indicated, maximum projections are shown. Image processing was performed using ImageJ software (Schindelin et al., 2012).

### Preparation of the cell derived matrix

To prepare the cell derived matrix, cells were grown for 3 days on coverslips until they reached confluence and then 50 ug/ml L-ascorbic acid-2-phosphate (Sigma Aldrich) was added to cultures. Cells were left for a further seven days. To prepare samples, cells were washed in PBS and extracted using pre-warmed extraction buffer (20 mM NH4OH, 0.5% Triton X-100 in PBS; 3 ml per 6 cm plate for 2 minutes). After three water washes, residual DNA was digested with 10 ug/ml DNase I (Roche) for 30 min at 37°C and then extracts were washed again. For immunofluorescence experiments, the matrix was fixed in 4% PFA and stained as above. For scanning electron microscopy, extracts were fixed with 2% PFA, 2.5% glutaraldehyde in 0.1M cacodylate buffer for 2 hours at room temperature. All coverslips were then washed 2x with 0.1M cacodylate buffer before undergoing progressive dehydration with 70, 80, 90, 96, 100, 100% EtOH for 15 minutes each. They were left in 100% ETOH for 1 h before drying in a critical point drier (50% spin, 120s delay between cycles, 15 CO2 cycles (13C cool, 15 cycles, 35C heat), speed 5). Samples were then sputter coated (40mA and 30 seconds). Coverslips were imaged on a FEI Quanta 200 FEG-SEM microscope.

### Live imaging

Procollagen trafficking was analysed using the retention using selective hooks (RUSH) assay (Boncompain et al., 2012). Cells stably expressing pro-SBP-GFP-COL1A1 were grown on 35 mm MatTek glass-bottomed dishes (MatTek) and transfected with Str-KDEL-IRES-ST-mCherry (McCaughey et al., 2019) 24 hours prior to imaging. Cells were always confluent at time of imaging.

To image, growth media was replaced with 1 ml prewarmed Fluorobrite DMEM (Thermo-Fisher, A1896701) and the dishes mounted on a Leica SP8 confocal laser scanning microscope system with a 63x HC OL APO CS2 1.42 numerical aperture glycerol lens and an environmental chamber at 37°C with CO2 enrichment. The microscope was controlled with Leica LAS X software. Once GFP- and mCh-positive cells were identified, 1 ml Fluorobrite containing 800 μM biotin and 1000 μg/ml L-ascorbic acid-2-phosphate (Sigma Aldrich, A92902) was added to the dish (final concentration 400 μM biotin and 500 μg/ml respectively) to induce procollagen folding and synchronous release of the pro-SBP-GFP-COL1A1 from the ER hook. This is T = 0. Cells were then imaged live, taking a single confocal plane image every 20 seconds. Fluorophores were excited using a 65-mW Argon and 20-mW solid state yellow lasers and detected using hybrid GaAsP detectors. Imaging conditions for movies: 1024×1024 pixel resolution, scanning speed 600 Hz, 2x zoom, 3-line average, imaging sequentially between frames. Fluorophores were excited using a 65-mW Argon laser at 488 nm and a 20-mW solid state yellow laser at 561 nm and detected using hybrid GaAsP detectors and corresponding notch filters.

To immediately fix during live imaging, 2 ml 8% PFA was added to the 2 ml Fluorobrite already present at the desired time point. Cells were fixed for 10 minutes and the above immunofluorescence protocol applied.

### Secretion assays, IPs and immunoblotting

To perform secretion assays, media was aspirated from confluent cells and replaced with a minimal volume of serum free DMEM-F12 supplemented with 50 μg/ul L-ascorbic acid-2-phosphate (Sigma Aldrich, A92902). Cells were left overnight at 37°C and 5% CO_2_. Media was then collected and the cell layer was rinsed with PBS, lysed in RIPA buffer (50 mM Tris-HCl, pH 7.5, 300 mM NaCl, 2% Triton X-100, 1% deoxycholate, 0.1% SDS, 1 mM EDTA) for 30 minutes rocking on ice, and collected without scraping. Media and lysate fractions were spun at 13000 x g at 4°C for 10 minutes and the supernatants used for SDS-PAGE. For collection of lysates in other experiments, cells were scraped in RIPA buffer, collected in tubes and incubated on a rotator at 4°C for 30 minutes. BCA assays were performed where necessary to determine protein concentration according to the manufacturer’s instructions (BioRad). Samples were boiled in 1x LDS sample buffer (Life Technologies, NP007) containing sample reducing agent (Life Technologies, NP007) for 10 minutes at 95°C.

For inhibitor experiments, confluent cells were treated with 1 ml serum free medium containing either DMSO (control), 10 μM MG132 or 200 nM bafilomycin overnight before media and lysate collection.

For GFP-trap^®^ immunoprecipitation, confluent cells were serum starved for 24 h in the presence of 50 μg/ul L-ascorbic acid-2-phosphate (Sigma Aldrich, A92902). At the time of assay, media was collected and spun at 2700 x g for 5 minutes to remove dead cells. The cell layer was washed twice with ice-cold PBS and lysed in ice cold buffer (10 mM Tris-HCl pH 7.4, 50 mM NaCl, 0.5 mM EDTA, 1.0% Igepal CA-630, 1 mM PMSF and 1x Protease inhibitor cocktail (Millipore, 539137)) for 30 minutes with agitation on a rotator. Lysate supernatant was collected by centrifuging at 20 000 x g for 10 minutes and 66 μl was removed to be used as an ‘input’ reference sample. Media and lysate supernatants were then incubated with equilibrated GFP nano-trap beads (Chromotek) on a rotator for two hours. Beads were pelleted by centrifugation at 2000 x g for 2 minutes. The supernatant was removed but 66 μl kept as an ‘unbound’ reference sample. The beads were then washed three times with 500 μl dilution buffer (10 mM Tris-HCl pH 7.4, 50 mM NaCl, 0.5 mM EDTA, 1 mM PMSF and 1x Protease inhibitor cocktail), pelleting them each time by centrifugation at 2000 x g for 2 minutes. All steps to this point were carried out on ice/at 4°C. Finally, beads were resuspended in 88 μl 1x LDS sample buffer (Life Technologies, NP007) containing sample reducing agent (Life Technologies) Inbound and unbound samples were mixed with 22 μl 4x LDS sample buffer + reducing agent (Life Technologies). Samples were boiled at 95 °C for 10 mins.

Secretion assays, IP and lysate samples were separated by SDS-PAGE on precast 3-8% tris-acetate and 4-12% bis-tris acrylamide gels (Invitrogen) followed by transfer to 0.2 μm nitrocellulose membranes (Amersham). Membranes were blocked in 5% milk 0.05% Tween (Sigma) in tris buffered saline for at least 1h. Primary and HRP-conjugated secondary antibodies were diluted in block and sequentially incubated with the membrane for 2-16 h each. HRP was detected by enhanced chemiluminescence (Promega ECL, GE healthcare film).

### Statistical analyses

Measurements of images and western blots were carried out using Image J. Statistical analyses were performed using GraphPad Prism 7.00. The tests used, n numbers and sample sizes are indicated in the figure legends, P-values where significant (p<0.05) are shown on the figures. All experiments were analysed with non-parametric tests as data was not normally distributed. Sample sizes were chosen based on previous, similar experimental outcomes and based on standard assumptions. No samples were excluded.

